# HIV-1 Nef interacts with LMP7 to attenuate immunoproteasome formation and MHC-I antigen presentation

**DOI:** 10.1101/697946

**Authors:** Yang Yang, Weiyong Liu, Dan Hu, Rui Su, Man Ji, Yuqing Huang, Muhammad Adnan Shereen, Xiaodi Xu, Zhen Luo, Qi Zhang, Fang Liu, Kailang Wu, Yingle Liu, Jianguo Wu

## Abstract

Proteasome is major protein degradation machinery and plays essential roles in diverse biological functions. Upon cytokine inductions, proteasome subunits β1, β2, and β5 are replaced by β1i/LMP2, β2i/MECL-1, and β5i/LMP7, leading to the formation of immunoproteasome. Immunoproteasome-degraded products are loaded onto the major histocompatibility complex class I (MHC-I) to regulate immune responses and induce cytotoxic-T-lymphocytes (CTLs). Human immunodeficiency virus type 1 (HIV-1) is the causal agent of acquired immunodeficiency syndrome (AIDS). HIV-1-specific CTLs represent critical immune responses to limit viral replication. HIV-1 negative regulatory factor (Nef) counteracts host immunity, especially the MHC-I/CTL. This study reveals a distinct mechanism by which Nef facilitates immune evasion through attenuating the functions of immunoproteasome and MHC-I. Nef interacts with LMP7 on the endoplasmic reticulum (ER) to down-regulate the incorporation of LMP7 into immunoproteasome, and thereby attenuating immunoproteasome formation. Moreover, Nef represses immunoproteasome protein degradation function, MHC-I trafficking, and antigen presentation activity.

**Importance:** Ubiquitin-proteasome system (UPS) is essential for degradation of damaged proteins, which takes place in proteasome. Upon cytokine inductions, proteasome catalytic activities are replaced by distinct isoforms resulting in formation of immunoproteasome. Immunoproteasome generates peptides for MHC-I antigen presentation and plays important roles in immune responses. HIV-1 is the agent of AIDS, and HIV-1-specific CTLs represent immune responses to limit viral replication. This study reveals a distinct mechanism by which HIV-1 promotes immune evasion. Viral protein Nef interacts with immunoproteasome component LMP7 to attenuate immunoproteasome formation and protein degradation function, and repress MHC-I antigen presentation activity. Therefore, HIV-1 targets LMP7 to inhibit immunoproteasome activation and LMP7 may be used as a target for the development of anti-HIV-1/AIDS therapy.

## Introduction

Proteasome is essential for protein degradations and carries out important processes such as clearance of mutated or misfolded proteins, cell signaling, and antigen presentation (1, 2). The 20S core particle of constitutive proteasome (cProteasome) is a barrel-shaped structure composed of 4 stacked rings: 2 outer rings contain α-subunits and act as binding sites for regulatory complexes to access proteins into proteasome inner chamber, and 2 inner rings contain β-subunits including β1, β2, and β5 active sites referred to “caspase-”, “trypsin-”, and “chymotrypsin-like” (3). Upon cytokine stress such as interferon (IFN) induction, the three catalytic subunits are replaced by distinct isoforms, β1i (LMP2), β2i (MECL-1), and β5i (LMP7), leading to formation of immunoproteasome (iProteasome) (4, 5). Full length low molecular mass protein 7 (proLMP7) is cleaved to a matured protein (mLMP7) for iProteasome assembly, and LMP7 is a key factor for iProteasome formation (6, 7). Mapping of LMP2 and LMP7 to major histocompatibility complex class I (MHC-I) locus combined with IFN-induced activation led to propose that iProteasome carries out a role in generating peptides for MHC-I antigen presentation (5). MHC-I is an efficient surveillance system to recognize and present antigens when hosts are invaded by pathogens. A critical step in the MHC-I pathway is processing antigens into smaller peptides and loading peptides onto a peptide loading complex (PLC) (8–10). MHC-I presents a diverse array of antigenic peptides (immunopeptidome) to circulate cytotoxic T lymphocytes (CTLs) (11, 12). Antigen processing pathway consists of sequential steps to ensure peptide-MHC-I assembly in the endoplasmic reticulum (ER) (13).

Viruses infect hosts, reproduce inside living cells, and must counteract and evade immune defense. Human immunodeficiency virus type 1 (HIV-1) is the causal agent of acquired immunodeficiency syndrome (AIDS) (14, 15). HIV-1-specific CTLs represent critical immune responses to limit viral replication (16–18) and MHC-I variants are determinants associated with AIDS disease progression (19, 20). HIV-1 genome encodes 5 proteins essential for viral replication and 4 accessory proteins (21, 22). The negative regulatory factor (Nef), an accessory protein of HIV-1, counteracts host immunity by interacting with PACS-1 and PI3K to attenuate MHC-I to the trans-Golgi network (23), down-regulates CD4 by hijacking AP-2 (24), enhances viral infectivity by excluding SERINC3/5 into the virions (25, 26), and induces secretion of exosomes from infected cells to stimulate viral spread (27).

This study reveals a distinct mechanism by which HIV-1 promotes immune evasion through attenuating the functions of immunoproteasome and MHC-I. Nef interacts with LMP7 on the ER membrane and down-regulates the incorporation of LMP7 into the iProteasome, and thereby attenuates iProteasome formation. Moreover, Nef represses iProteasome protein degradation function and inhibits MHC-I trafficking and antigen presentation activity.

## Results

### LMP7 is associated with Nef

Nef is a critical protein necessary for HIV-1 pathogenesis, immune evasion, and viral spread. To evaluate the mechanism by which Nef regulates immune response, we initially utilized a yeast two-hybrid system to screen proteins interacting with Nef (Fig. 1A). The results showed that LMP7 was associated with Nef. Co-immunoprecipitation (Co-IP) indicated that Nef interacted with LMP7 in human embryonic kidney (HEK) 293T cells (Fig. 1B). Protein-protein pull-down revealed that purified Nef-GST directly interacted with LMP7 (Fig. 1C). A yeast two-hybrid experiment further showed that Nef associated with LMP7 (Fig. 1D). Moreover, the interacting strength between LMP7 and Nef was determined by BioLayer Interferometry (BLI). Association of LMP7 with Nef was robust, wavelength shift value of LMP7 and Nef-GST (71.1 pm) was significantly higher as compared to that of LMP7 and GST (0.7 pm) (Fig. 1E), confirming that LMP7 is tightly associated with Nef. Meanwhile, a series of concentrations of Nef protein were tested with the same amount of LMP7 protein, the wavelength shift values were increased as higher concentration of Nef tested, and binding affinity of LMP7 with Nef measured by Kd value was high (734.7 nM) (Fig. 1F).

**Figure 1.**
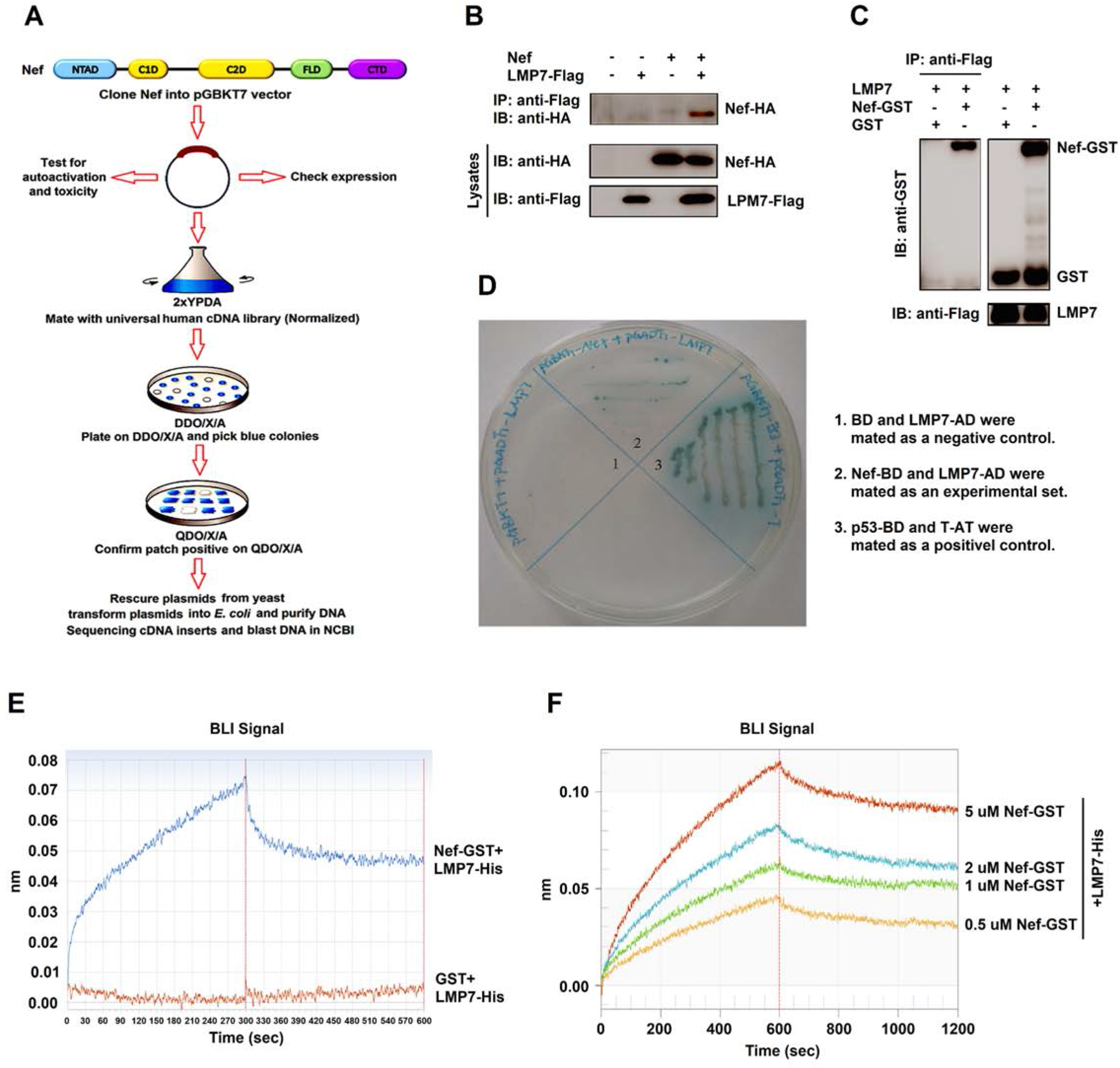
LMP7 interacts with Nef. (**A**) Flow chart of yeast two-hybrid system to screen proteins interacting with Nef. (**B**) 293T cells were transfected with pNef-HA and pLMP7-Flag for 24 h and cell lysates were prepared for co-immunoprecipitation (CoIP). Anti-Flag antibodies were used for CoIP and anti-HA antibodies were used to detect Nef-HA. Cell lysates were detected with anti-HA and anti-Flag antibodies. (**C**) Purified LMP7-His-Flag was mixed with GST or Nef-GST. Proteins were pulled down by anti-Flag antibodies (IP), and detected with anti-GST antibodies (IB). Cell lysates were detected with anti-GST and anti-Flag antibodies. (**D**) Yeast two-hybrid system to conform interaction between Nef and LMP7. Mated yeast cells of Nef fused with Gal4 DNA-binding domain (BD) and LMP7 fused with Gal4 activation domain (AD). BD and LMP7-AD were mated as a negative control by using agar containing QDO (-Ade/-His/-Leu/-Trp) and X-α-Gal (Area 1). Nef-BD and LMP7-AD were mated as an experimental set (Area 2). P53-BD and T-AD were mated as a positive control (Area 3). (**E**) Biolayer interferometry (BLI)wave length shifts of Nef-GST with LMP7-His (blue) and GST with LMP7-His (red) were recorded during the processes of association and dissociation. (**F**) Biolayer interferometry (BLI)wave length shifts of increasing concentrations of Nef-GST with the same amount of LMP7-His were recorded during the processes of association and dissociation. All the results are the representatives of three independent experiments.

### LMP7 binds to Nef through the sequences from amino acids 69 to 160

It is known that full length LMP7 (proLMP7) is cleaved into a matured LMP7 (mLMP7) for the iProteasome assembly upon IFN-γ induction. To evaluate the requirement of the two forms of LMP7 in interacting with Nef, plasmids expressing proLMP7 and mLMP7 were constructed (Fig. 2A). Both proLMP7 and mLMP7 were able to pull down Nef (Fig. 2B), indicating that mLMP7 is sufficient for the interaction, although the full length of LMP7 was able to pull down more Nef than mLMP7. Three fusion proteins were generated, in which the green fluorescent protein (GFP) was fused to proLMP7, mLMP7, and pro-domain of LMP7 (Fig. 2C). proLMP7, proLMP7-GFP, and mLMP7-GFP were able to pull down Nef, but proD-GFP failed to act (Fig. 2D), suggesting that pro-domain is not involved in interacting with Nef. The GFP fused proLMP7 and mLMP7 pulled down comparable amount of Nef, however both of them were able to pull down more Nef than proLMP7 (Fig. 2D). To narrow down the sequences interacting with LMP7, 4 deletions of LMP7 were generated (Fig. 2E). Nef interacted with LMP7 and LMP7(69–272) strongly, with LMP7(1-260) and LMP7(1-240) moderately, and with LMP7(1-200) and LMP7(1-160) weakly (Fig. 2F), indicating that amino acids from 69 to 160 of LMP7 are required for the interaction with Nef. Furtherly, we tested an LMP7 truncation without the domain from 69 to 160 residues (Fig. 2G), which showed that this truncation proD/Tail failed to interact with Nef, implying that this region is crucial for the binding with Nef (Fig. 2H).

**Figure 2.**
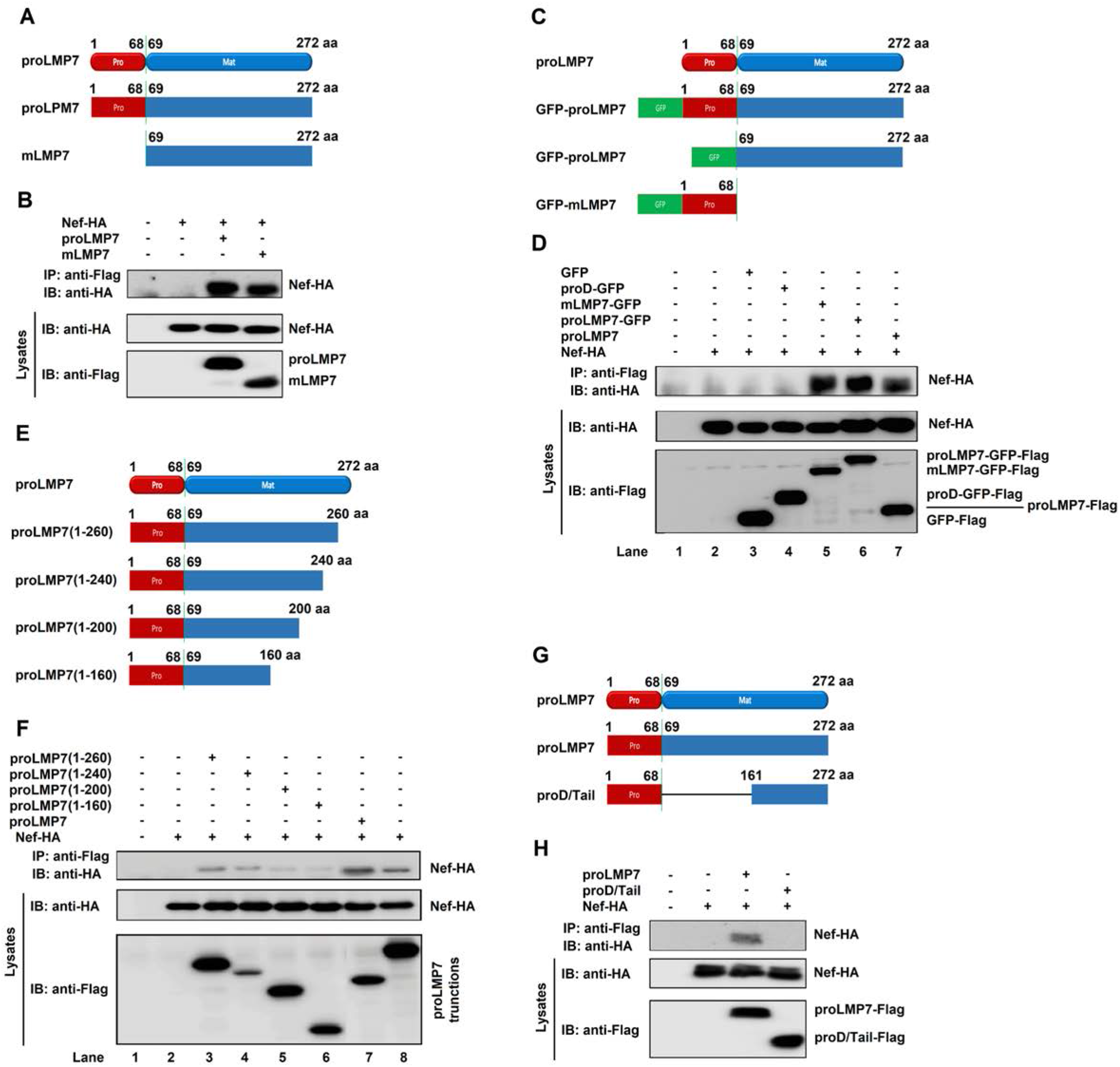
LMP7 interacts with Nef by sequences from amino acids 69 to 160. (**A**) Schematic structures of proLMP7 and mLMP7. (**B**) 293T cells were co-transfected with pNef-HA and pproLMP7-Flag or pmLMP7-Flag for 24 h and cell lysates were prepared for Co-IP. (**C**) Schematic structures of fusion proteins, proLMP7-GFP, mLMP7-GFP, and proD-GFP. (**D**) 293T cells were co-transfected with p-Nef-HA and p-GFP-Flag, p-proD-GFP-Flag, p-mLMP7-GFP-Flag, p-proLMP7-GFP-Flag, or p-proLMP7-Flag for 24 h and cell lysates were prepared for CoIP. (**E**) Schematic structures of truncated LMP7. (**F**) 293T cells were co-transfected with pNef-HA and pLMP7 (1-260)-Flag, pLMP7 (1-240)-Flag, pLMP7 (1-200)-Flag, pLMP7 (1-160)-Flag, pLMP7 (69-272)-Flag, or pLMP7-Flag for 24 h and cell lysates were prepared for CoIP. (**G**) Schematic structures of proLMP7 and proD/Tail. (**H**) 293T cells were co-transfected with pNef-HA and pproLMP7-Flag or proD/Tail-Flag for 24 h and cell lysates were prepared for Co-IP. (**B**, **D**, **F, H**) Anti-Flag antibodies were used for IP, and detected with anti-HA antibodies. Cell lysates were detected with anti-HA and anti-Flag antibodies. All the results are the representatives of three independent experiments.

### Nef interacts with LMP7 by the C1D and C2D domains

The domains of Nef interacting with LMP7 were also determined. 4 truncated mutants of Nef were constructed (Fig. 3A). LMP7 interacted with Nef, Nef(1-149), and Nef(85-206), but failed to interact with Nef(1-84) and Nef(150-206) (Fig. 3B), suggesting that amino acids from 85 to 149 of Nef are essential for the interaction. Next, 3 truncations within amino acids 85 to 149 were constructed (Fig. 3C). LMP7 interacted with Nef(1-115), Nef(1-130), and Nef(1-149), but not with Nef(1-84) or Nef(1-100) (Fig. 3D), demonstrating that residues from 101 to 115 of Nef are required for interacting with LMP7. We also observed that the interacting strength between Nef truncations and LMP7 was gradually increased, Nef(1-115) was the weakest, Nef(1-149) was the strongest, and Nef(1-130) was modest (Fig. 3D). We further determined which residues of Nef were essential for interacting with LMP7 by generating four site-directed mutations (Fig. 3E). LMP7 strongly interacted with Nef(1-115) and Nef(1-115)-(105-108A), weakly interacted with Nef (1-115)-(101-104A), but not interacted with GFP, Nef(1-115)-(109-112A), or Nef(1-115)-(113-115A) (Fig. 3F), indicating that residues from 109 to 115 are crucial for the interaction. The 7 residues were individually mutated to Alanine in Nef(1-115) (Fig. 3G). LMP7 strongly interacted with Nef(1-115), Nef(1-115)-(I109A), Nef(1-115)-(L110A), Nef(1-115)-(D111A), Nef(1-115)-(W113A), and Nef(1-115)-(I114A), weakly interacted with Nef(1-115)-(L112A), and barely interacted with Nef(1-115)-(Y115A) (Fig. 3H), suggesting that residues 12 and 15 are required for the interaction. Finally, a double residue mutant was generated to confirm the role of L112 and Y115 in the interaction with LMP7 (Fig. 3I). LMP7 interacted with Nef(1-1. 110) and Nef(1-115), weakly interacted with Nef(1-115)-(L112A), barely interacted with Nef(1-115)-(Y115A), but not interacted with Nef(1-115)-(L112A/Y115A) (Fig. 3J), demonstrating that L112 and Y115 within the C2D domain of Nef are essential for the interaction. Several amino acid residues within the C2D domain are essential for Nef oligomerization and interaction with host proteins (28–30). We demonstrate that residues L112 and Y115 within this domain are required for interacting with LMP7.

**Figure 3.**
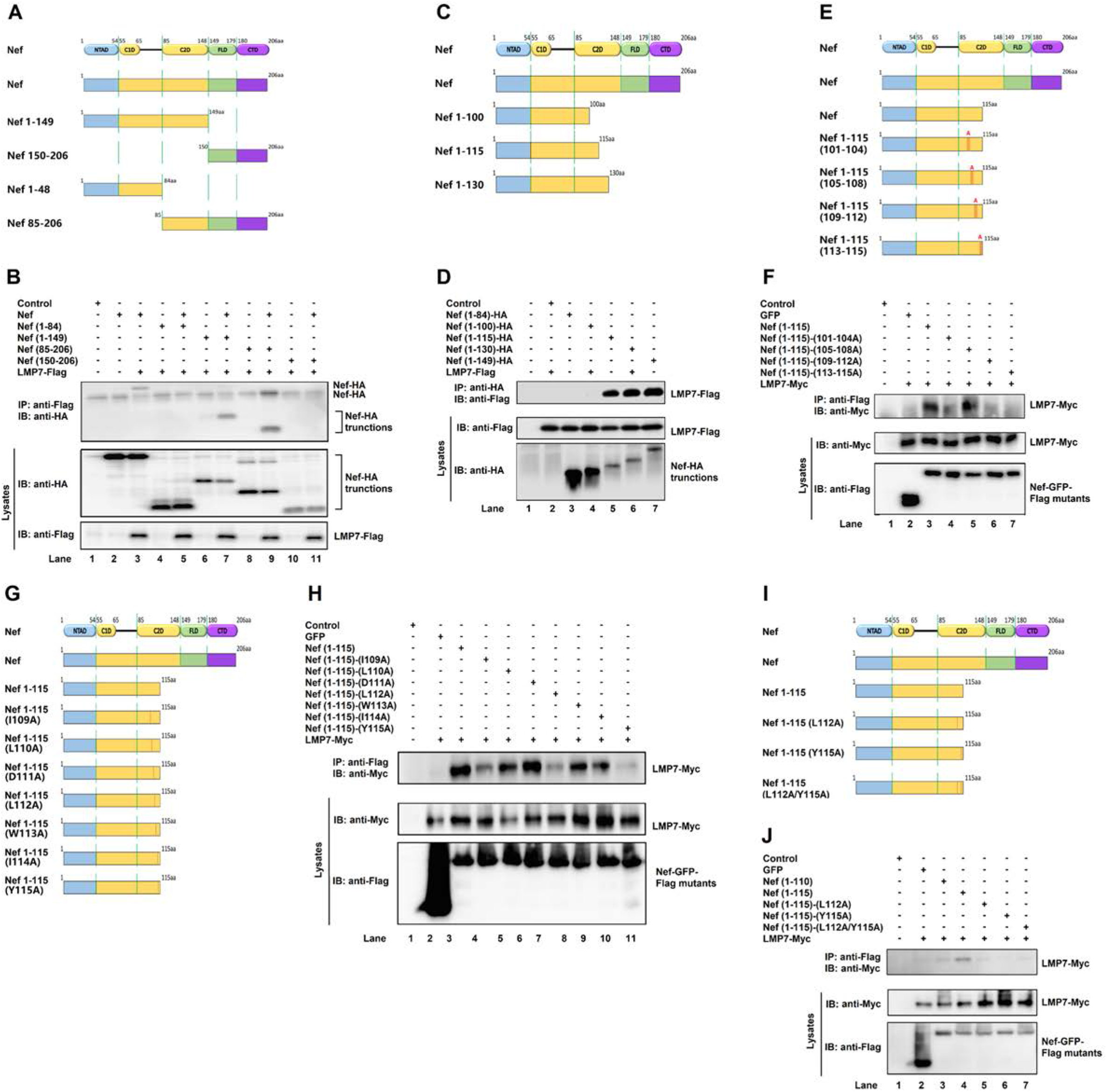
Nef interacts with LMP7 through the C1D and C2D domains. (**A**) Schematic structures of Nef and truncated Nef. NTAD: N-terminal anchor domain; C1D: core 1 domain; C2D: core 2 domain; FLD: flexible loop domain; CTD: C-terminal domain. (**B**) 293T cells were co-transfected with pLMP7-Flag and pNef-HA, pNef (1-84)-HA, pNef (1-149)-HA, pNef (85-206)-HA, or pNef (150-206)-HA for 24 h. (**C**) Schematic structures of Nef truncations within amino acids 85 to 149. (**D**) 293T cells were co-transfected with pLMP7-Flag and pNef (1-84)-HA, pNef (1-100)-HA, pNef (1-115)-HA, pNef (1-130)-HA, or pNef (1-149)-HA for 24 h. (**B**, Cell lysates were prepared for Co-IP. Anti-Flag antibodies were used for IP and detected with anti-HA antibodies. Cell lysates were detected with anti-HA and anti-Flag antibodies. (**E**) Schematic structures of Nef point mutations, in which 3 or 4 residues in Nef (1-115) were replaced by alanine. (**F**) 293T cells were co-transfected with pLMP7-Myc and pGFP-Flag, pNef (1-115)-Flag, pNef (1-115)-(101-104A)-Flag, pNef (1-115)-(105-108A)-Flag, pNef (1-115)-(109-112A)-Flag, or pNef (1-115)-(113-115A)-Flag for 24 h. (**G**) Schematic structures of 7 Nef single residue mutations, in which residues were mutated individually to alanine in Nef (1-115). (**H**) 293T cells were co-transfected with pLMP7-Myc and pGFP-Flag, pNef (1-115)-Flag, pNef (1-115)-(I109A)-Flag, pNef (1-115)-(L110A)-Flag, pNef (1-115)-(D111A)-Flag, pNef (1-115)-(L112A)-Flag, pNef (1-115)-(W113A)-Flag, pNef (1-115)-(I114A)-Flag, or pNef (1-115)-(Y115A)-Flag for 24 h. (**I**) Schematic structure of 2 Nef single mutants and 1 Nef double mutant. (**J**) 293T cells were co-transfected with pLMP7-Myc and pGFP-Flag, pNef (1-110)-Flag, pNef (1-115)-Flag, pNef (1-115)-(L112A)-Flag, pNef (1-115)-(Y115A)-Flag, or pNef (1-115)-(L112A/Y115A)-Flag for 24 h. (**F**, **H**, **J**) Cell lysates were prepared for Co-IP. Anti-Flag antibodies were used for IP and detected with anti-Myc antibodies. Cell lysates were detected with anti-Myc and anti-Flag antibodies. All the results are the representatives of three independent experiments.

### Nef interacts with LMP7 to attenuate immunoproteasome formation

The 20S core particle of constitutive proteasome (cProteasome) is a barrel-shaped structure composed of 4 stacked rings: 2 outer rings contain α-subunits and act as binding sites for regulatory complexes to access proteins into proteasome inner chamber, and 2 inner rings contain β-subunits including β1, β2, and β5 active sites referred to “caspase-”, “trypsin-”, and “chymotrypsin-like” (3). Upon cytokine stress such as interferon (IFN) induction, the three catalytic subunits are replaced by distinct isoforms, β1i (LMP2), β2i (MECL-1), and β5i (LMP7), leading to formation of immunoproteasome (iProteasome) (4, 5). The biological effect of the interaction between Nef and LMP7 was evaluated. LMP7 is a key factor for iProteasome assembly (Fig. 4A, B) (31–33). It is reasonable to speculate that Nef may affect iProteasome formation by interaction with LMP7. To verify this, HeLa cells were treated with recombinant human IFN-γ (rhIFN-γ). proLMP7 was expressed in the absence of rhIFN-γ, but not detected in the presence of rhIFN-γ; in contrast, mLMP7 was not detected in the absence of rhIFN-γ, but detected in the presence of rhIFN-γ (Fig. 4C); suggesting that IFN-γ induces LMP7 maturation. Cell lysates, proteasomes, and iProteasomes were prepared from the cells transfected with Nef and treated with rhIFN-γ. β-actin was not detected in proteasomes or iProteasomes, but detected in cell lysates (Fig. 4D), revealing that the proteasomes were successfully purified (34). Endogenous LMP7 was not detected in proteasomes and cell lysates in the absence of rhIFN-γ, but detected in iProteasomes and cell lysates in the presence of rhIFN-γ (Fig. 4D). In the presence of rhIFN-γ, Nef was not detected in iProteasome, but detected in cell lysate (Fig. 4D), suggesting that Nef is not associated with fully assembled iProteasome. Interestingly, LMP7 was significantly attenuated by Nef in iProteasome, but relatively unaffected by Nef in cell lysate (Fig. 4D). Moreover, LMP7 in iProteasomes was reduced by 80% in the presence of Nef (Fig. 4E), suggesting that Nef attenuates the incorporation of LMP7 into iProteasome and thereby down-regulates iProteasome formation. To further investigate whether myristoylation of Nef would affect the formation of iProteasomes, the second amino acid glycine on Nef, which is critical for the myristoylation of Nef (35), was mutated to alanine as Nef-G2A. Intriguingly, LMP7 in iProteasomes was much decreased when Nef or Nef-G2A expressed, compared to samples that empty vector transfected, when the cells were treated with rhIFN-γ, even the expressions of LMP7 in both Nef and Nef-G2A transfected cells were higher than in empty vector transfected cells in cell lysates (Fig. 4F). Meanwhile, both of Nef and Nef-G2A were able to reduce LMP7 in iProteasomes to 20% in the presence of rhIFN-γ (Fig. 4G), indicating that Nef myristoylation may be not related to the formation of iProteasome. In order to study the effect of the association of Nef and LMP7 on the assembly of iProteasomes, 3 Nef mutants were tested. LMP7 was highly attenuated by Nef(1–115) in iProteasomes, whereas was less attenuated by both Nef(1-115)-(109-112A) and Nef(1-115)-(103-115A) (Fig. 4H). Meanwhile, LMP7 in iProteasomes was reduced by about 90% in the presence of Nef, whereas 40% in the presence of Nef(1-115)-(109-112A) and 20% in the presence of Nef(1-115)-(103-115A) (Fig. 4I), suggesting that residues from 109 to 115 on Nef could be crucial for the inhibition of iProteasome formation.

**Figure 4.**
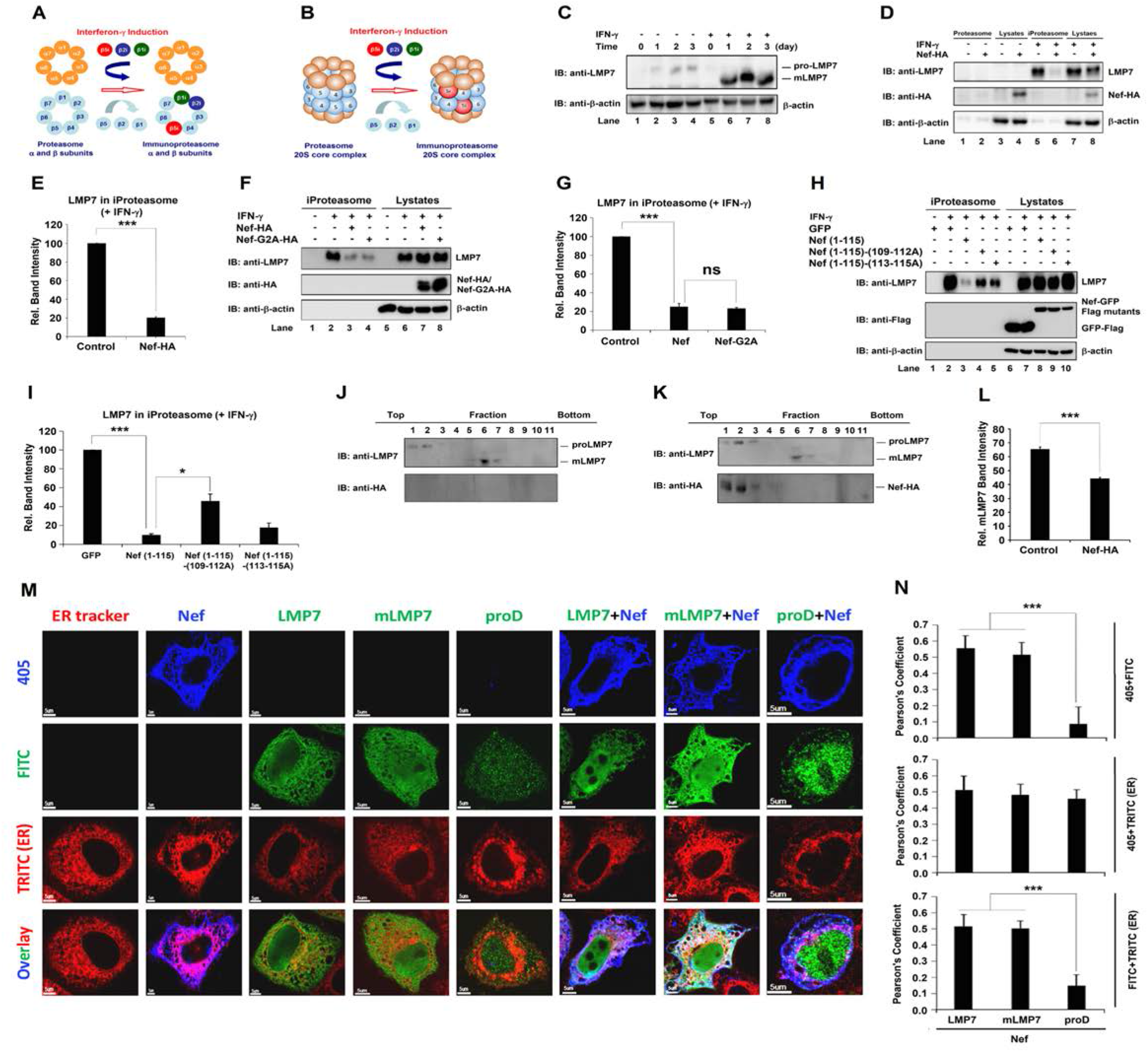
Nef interacts with LMP7 to attenuate immunoproteasome formation. (**A**) Schematic structures of α and β rings of cProteasome and iProteasome. Upon IFN-γ stimulation, immunoproteasome specific subunits β1i, β2i, and β5i were assembled into iProteasome. (**B**) Schematic structures of 20S core particles of proteasome and iProteasome. Upon IFN-γ stimulation, immunoproteasome specific subunits β1i, β2i, and β5i were assembled into iProteasome. (**C**) HeLa cells were treated with rhIFN-γ for 0, 1, 2, and 3 days. LMP7 and β-actin in the cells were determined by Western blots. (**D**, **E**) Lysates, proteasomes, and iProteasomes were prepared from HeLa cells transfected with empty vector or pNef-HA for 24 h, followed by rhIFN-γ treatment or nontreatment for 24 h. LMP7, Nef, and β-actin were detected by Western blots (D). Relative band intensity stands for the ratio of LMP7 in immunoproteasomes to LMP7 in lysates. The relative band intensity of the empty vector was set to 100% as a negative control (E). (**F**, **G**) Lysates, proteasomes, and iProteasomes were prepared from HeLa cells transfected with empty vector, pNef-HA, or pNef-G2A-HA for 24 h, followed by rhIFN-γ treatment or nontreatment for 24 h. LMP7, Nef, Nef-G2A and β-actin were detected by Western blots (F). Relative band intensity stands for the ratio of LMP7 in immunoproteasomes to LMP7 in lysates. The relative band intensity of the empty vector was set to 100% as a negative control (G). (**H, I**) Lysates, proteasomes, and iProteasomes were prepared from HeLa cells transfected with pGFP-Flag, pNef (1-115)-Flag, pNef (1-115)-(109-112A)-Flag, or pNef (1-115)-(113-115A)-Flag for 24 h, followed by rhIFN-γ treatment or nontreatment for 24 h. LMP7, GFP, Nef (1-115), Nef (1-115)-(109-112A), or Nef (1-115)-(113-115A), and β-actin were detected by Western blots (H). Relative band intensity stands for the ratio of LMP7 in immunoproteasomes to LMP7 in lysates. The relative band intensity of pGFP-Flag was set to 100% as a negative control (I). (**J, K, L**) HeLa cells were transfected with empty vector (F) or pNef-HA (G) for 24 h, followed by rhIFN-γ treatment or nontreatment for 24 h, and cell lysates were collected as 11 fractions from top to bottom after ultracentrifugation. proLMP7, mLMP7, and Nef were determined by Western blots. Relative mLMP7 band intensity stands for the ratio of mLMP7 in all fractions to proLMP7 plus mLMP7 in all fractions. The relative band intensity of empty vector was set as a negative control (L). (**M, N**) HeLa cells were transfected with empty vector, pNef-HA, pLMP7-Flag, pmLMP7-Flag, pproD-Flag, pNef-HA+pLMP7-Flag, pNef-HA+pmLMP7-Flag, or pNef-HA+pproD-Flag for 24 h. Transfected cells were stained with ER tracker (red), anti-HA antibodies (blue) or anti-Flag antibodies (green). Pictures were taken using FluoView FV1000 (Olympus) confocal microscope. Pearson’s coefficient values were calculated by using the software OLYMPUS FLUOVIEW Ver.1.7a Viewer (N). All the results are the representatives of three independent experiments, and the average is calculated from three independent experiments and error bars are indicated as standard deviation of mean. NS, not significant, *P* ≤0.05 (*), *P* ≤0.01 (**), *P* ≤0.001 (***).

The role of Nef in regulation of iProteasome was further determined using gradient ultracentrifugation analyses. In the absence of Nef, proLMP7 was detected in the top fractions 1– 2, while mLMP7 was distributed in the bottom fractions 6–7 (Fig. 4J). In the presence of Nef, proLMP7 was detected in the top fractions 1–3, mLMP7 was presence in the bottom fractions 5– 8, and Nef was distributed in the top fractions 6–7 (Fig. 4K). These results further suggest that Nef is not associated with assembled iProteasome, but instead Nef interacts with LMP7 to prevent incorporation of LMP7 into iProteasome. Nef also exhibited the ability to inhibit LMP7 incorporated into iProteasome (Fig. 4L), which is similar as the data in Fig 4D. It was reported that critical steps of iProteasome formation occur on the ER membrane (36). We showed that both Nef and LMP7 were distributed mostly on the ER membrane in the cytosol, when they were expressed alone. While a large proportion of Nef and LMP7 also co-localized and distributed on the ER membrane when they were expressed together (Fig. 4M, N). Interestingly, Nef and mLMP7 co-localized and distributed mostly on the ER membrane in the cytosol too. However, Nef did not co-localized with proD, and Nef but not proD localized on the ER membrane in the cytosol, suggesting that the interaction between Nef and LMP7 plays an important role on the co-localization on the ER membrane (Fig. 4M, N). Thus, we demonstrate that a relatively large amount of Nef interacts with LMP7 on the ER membrane to prevent LMP7 from incorporation into iProteasome, and thereby attenuate iProteasome formation at early stage.

### Nef attenuates immunoproteasome protein degradation activity and MHC-I antigen presentation function

The central task of iProteasome is to degrade ubiquitylated proteins that are executed ubiquitin proteasome system (UPS) (Fig. 5A) (37, 38). The effects of Nef on ubiquitylated protein degradation function of iProteasome were determined. Cells were transfected with pNef and treated with rhIFN-γ. Ubiquitinated proteins were induced by rhIFN-γ and further enhanced by Nef (Fig. 5B, C). Cells were then infected with HIV-1 (pNL4-3) or Nef-deficient HIV-1 (pNL4-3dNef) and treated with rhIFN-γ. Ubiquitinated proteins were induced by rhIFN-γ, further enhanced by pNL4-3, but not affected by pNL4-3dNef (Fig. 5D, E). The results demonstrate that Nef protects protein ubiquitination by attenuating iProteasome protein degrading function.

**Figure 5.**
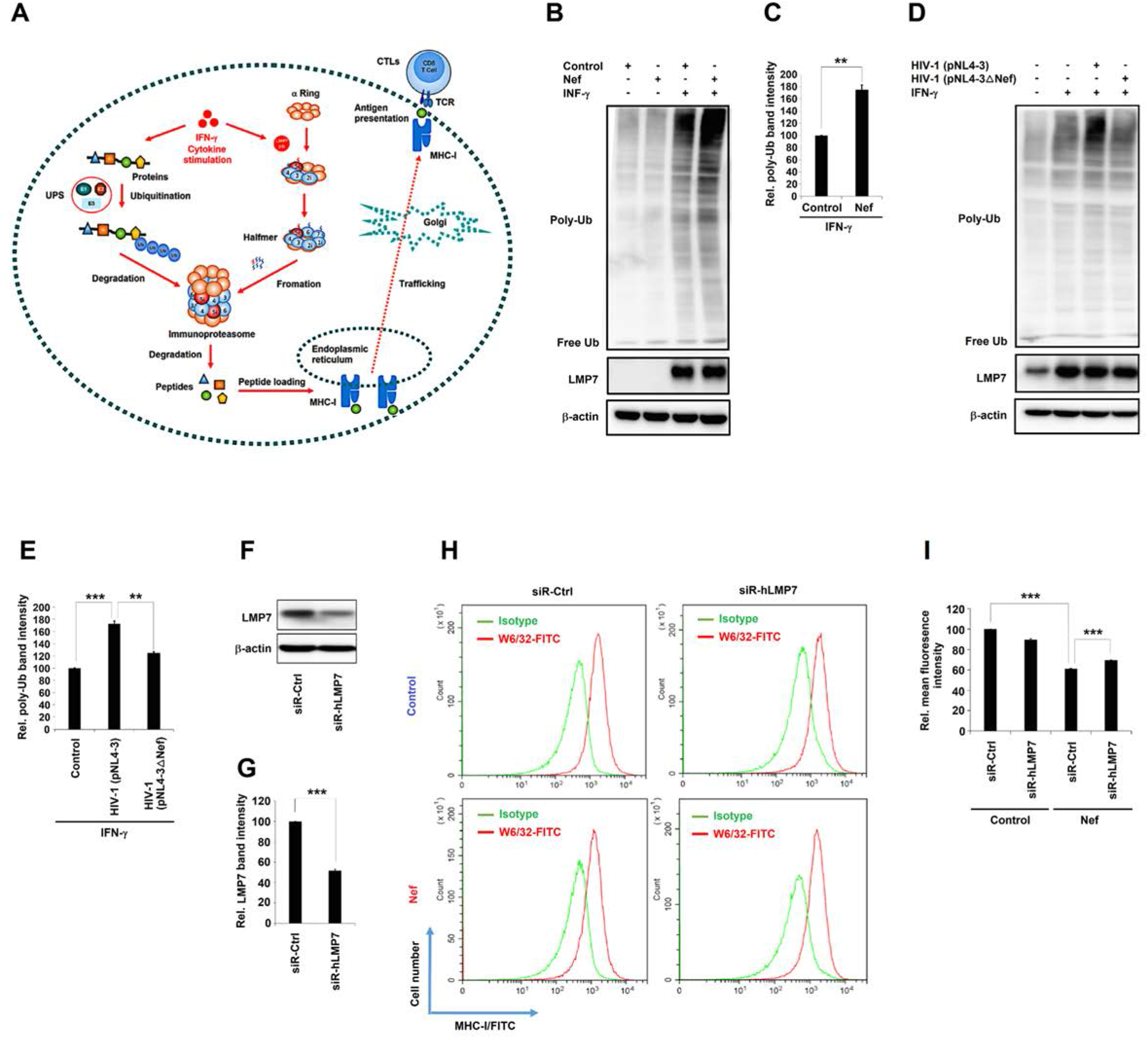
Nef attenuates immunoproteasome protein degradation activity and MHC-I antigen presentation function. Diagram of UPS, iProteasome, MHC-I, and CTL. (**B**, **C**) HeLa cells were transfected with pNef for 24 h and treated or untreated with rhIFN-γ. Ubiquitination of proteins and expression of LMP7 and β-actin were determined by Western blots (B). Relative poly-ub band intensity stands for the ratio of the ubiquitination of proteins to the correlated β-actin. The relative poly-ub band intensity of empty vector treated with rhIFN-γ was set to 100% as a negative control (C). (**D**, **E**) CD4^+^ Jurkat T cells were infected with HIV-1 (pNL4-3) or Nef-deficient HIV-1 (pNL4-3dNef) at MOI 10 ng P24/106 cells and treated with rhIFN-γ. Ubiquitination of proteins and expression of LMP7 and β-actin were determined by Western blots (D). Relative poly-ub band intensity stands for the ratio of the ubiquitination of proteins to the correlated β-actin. The relative poly-ub band intensity of uninfected cells treated with rhIFN-γ was set to 100% as a negative control (E). (**F, G**) CD4^+^ Jurkat T cells were transfected with siR-hCtrl or siR-hLMP7 for 48 h. Cell lysates were prepared and LMP7 and β-actin were detected by Western blots (F) and intensities of protein bands were quantified, and relative LMP7 band intensity stands for the ratio of LMP7 to β-actin band intensity (G). (**H, I**) pLenti-Flag or pLenti-Nef-Flag CD4+ Jurkat T cells were transfected with siR-hCtrl or siR-hLMP7 for 48 h. Cells were collected and accessed to flow cytometry with W6/32 anti-MHC I-FITC antibodies. Cells stained with IgG isotype antibody were served as MHC-I negative controls (H). Mean fluorescence intensities (MFI) were calculated for all W6/32 MHC I-FITC stained cells (I). All the results are the representatives of three independent experiments, and the average is calculated from three independent experiments and error bars are indicated as standard deviation of mean. NS, not significant, *P* ≤0.05 (*), *P* ≤0.01 (**), *P* ≤0.001 (***).

Immunoproteasomes process cellular proteins into peptides that are loaded onto MHC-I to communicate intracellular protein composition to immune system (39, 40). Deletion of LMP7 reduces MHC-I cell surface expression (41). Here, the effects of Nef on regulating MHC-I trafficking and antigen presentation were determined by using an siRNA, which is specifically targeting human LMP7 mRNA. In CD4+ Jurkat T cells, the expression of LMP7 was reduced by 50% in the presence of siR-hLMP7, compared to in the presence of siR-hCtrl (Fig. 5F, G). CD4+ Jurkat T cells were transfected with siR-hLMP7 or siR-hCtrl, which were negative controls, while cells stained with IgG isotype antibody were served as MHC-I negative controls (Fig. 5H). Additionally, the relative mean fluorescence intensity (MFI) was significantly reduced to 60% by Nef in the presence of siR-hCtrl (Fig. 5I), indicating that Nef attenuates MHC-I cell surface expression. Then the relative MFI was significantly up-regulated to 70% by Nef in the presence of siR-hLMP7 (Fig. 5I), suggesting that LMP7 is involved in Nef-mediated attenuation of MHC-I cell surface expression. Interestingly, the relative MFI was reduced to 90% in the presence of siR-hLMP7 (Fig. 5I), revealing that MHC-I trafficking is attenuated by siR-hLMP7. Therefore, the results demonstrate that MHC-I trafficking is repressed by Nef and attenuated by siR-hLMP7, and thus LMP7 is involved in Nef-mediated attenuation of MHC-I trafficking.

### Nef inhibits MHC-I antigen presentation pathway by hijacking LMP7

The iProteasome mediates immune responses by efficiently generating peptides for antigen presentation of MHC-I, which is an efficient surveillance system that present antigens when hosts are invaded by pathogens. The beginning step in the MHC-I signaling pathway is processing the antigenic proteins into peptides and loading them onto peptide loading complex (PLC), in which iProteasomes play a vital role. MHC-I molecules present a diverse array of antigenic peptides (immunopeptidome) to circulating TCLs. Here, the biological effect of Nef on MHC-I function was determined using an antigen-presenting cell (APC) system (42, 43). Ovalbumin (OVA) specific antigen presentation assay was carried out, in which mouse leukaemic monocyte macrophages (RAW264.7) were transfected with mouse siRNA siR-mLMP7 or control siRNA siR-mCtrl (Fig. 6A). The expression of LMP7 was reduced to 52% in the presence of siR-mLMP7, compared to in the presence of siR-mCtrl (Fig. 6B). Then RAW264.7 cells were transfected with pNef-Flag or empty vector and incubated with OVA, and the cells were co-cultured with mouse CD8^+^ T B3Z cells that recognize OVA-generated peptide and promote IL-2 secretion, an indicator of antigen presentation (Fig. 6C). Secreted IL-2 protein was induced by OVA in the presence of siR-mCtrl (from 2 pg/ml to 99 pg/ml), or in the presence of siR-mLMP7 (from 3 pg/ml to 81 pg/ml). However induction of IL-2 was inhibited by Nef in the presence of siR-mCtrl (from 99 pg/ml to 51 pg/ml), while this inhibition by Nef was significantly impaired in the presence of siR-mLMP7 (from 51 pg/ml to 65 pg/ml) (Fig. 6D, E), suggesting that Nef represses MHC-I antigen presentation and LMP7 is involved in such repression. Taken together, this study reveals a distinct mechanism by which HIV-1 Nef interacts with LMP7 to promote immune evasion by suppressing iProteasome formation and protein degradation function, and repressing MHC-I trafficking and antigen-presentation activity (Fig. 7).

**Figure 6.**
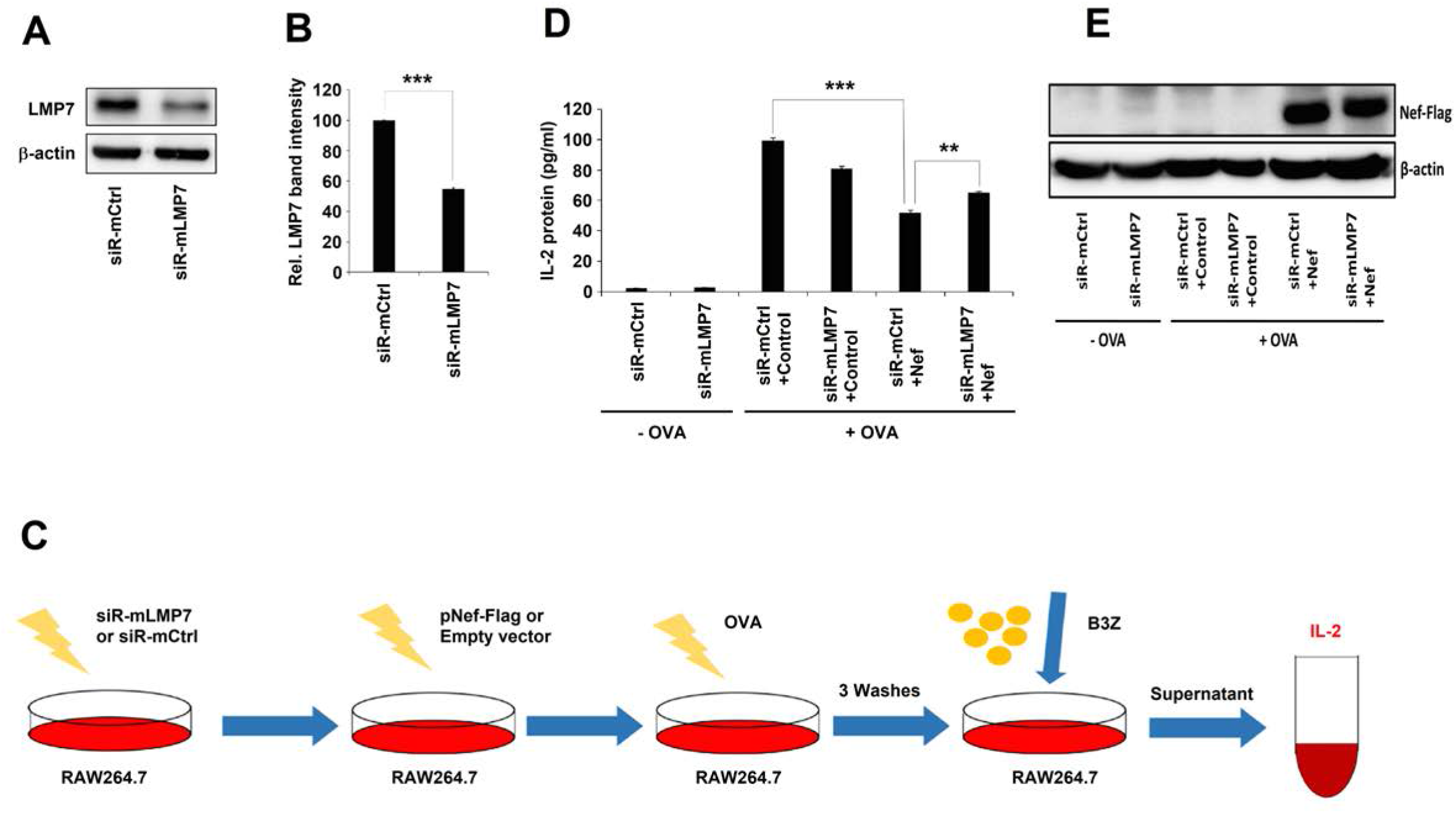
Nef inhibits MHC-I antigen presentation pathway by hijacking LMP7. (**A**, **B**) RAW 264.7 cells were transfected with siR-mCtrl or siR-mLMP7 for 48 h. Cell lysates were prepared and LMP7 and β-actin were detected by Western blots (A) and intensities of protein bands were quantified, and relative LMP7 band intensity stands for the ratio of LMP7 to β-actin band intensity (B). (**C**) Flow chat of ovalbumin (OVA) specific antigen presentation assay. (**D**, **E**) RAW264.7 cells were transfected with siR-mCtrl or siR-mLMP7 for 24 h, and then transfected with empty vector or pNef-Flag for 24 h. IL-2 secreted from B3Z cells were measured using ELISA. Cells untreated with OVA were set as negative controls (D). The expressions of Nef-Flag and β-actin were detected by Western blots (E). All the results are the representatives of three independent experiments, and the average is calculated from three independent experiments and error bars are indicated as standard deviation of mean. NS, not significant, *P* ≤0.05 (*), *P* ≤0.01 (**), *P* ≤0.001 (***).

**Figure 7.**
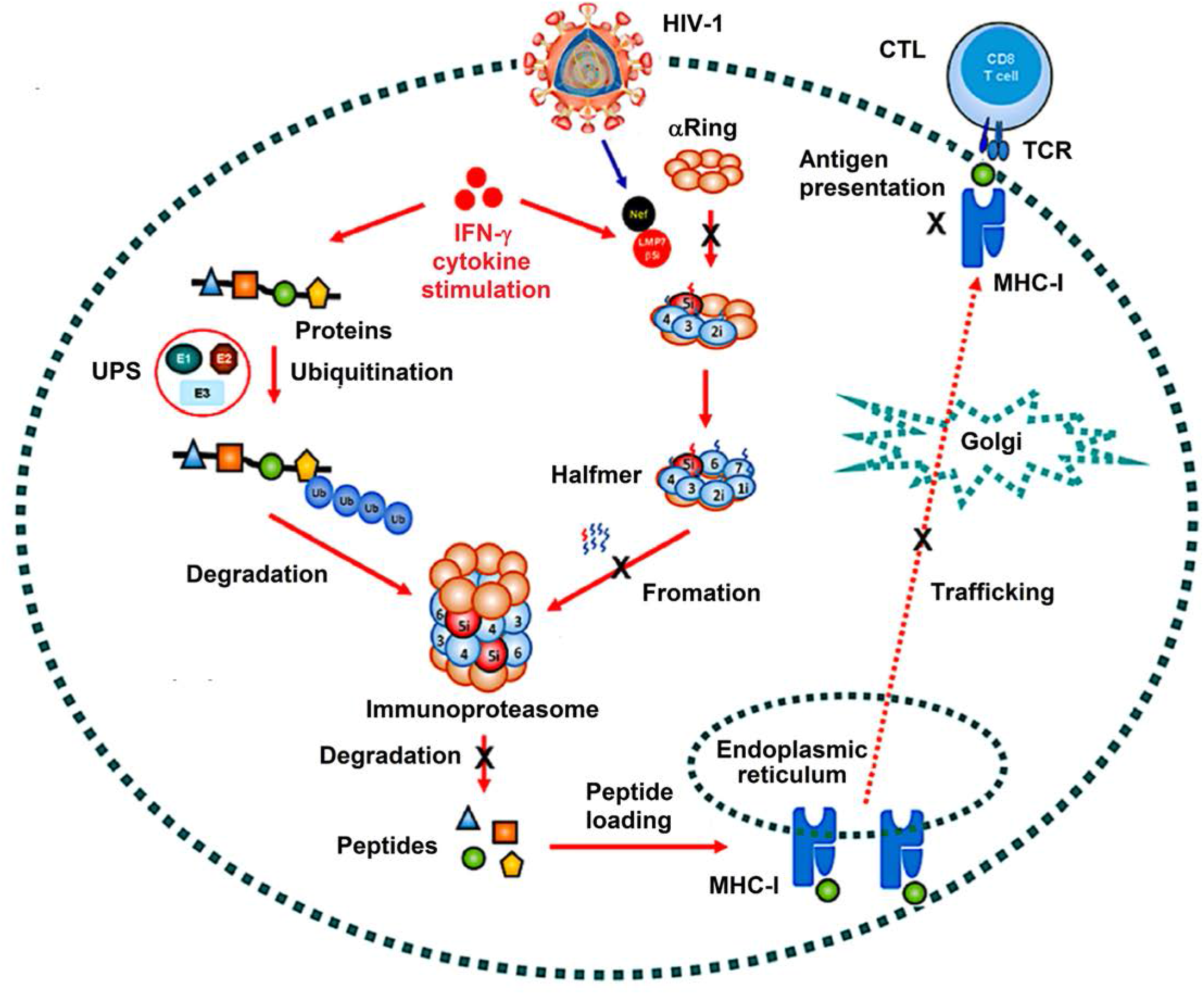
A proposed mechanism by which Nef represses the functions of immunoproteasome and MHC-I. The ubiquitin proteasome system (UPS) is the predominant system responsible for the degradation of 80% of all cellular proteins. Proteins are targeted for proteasomal degradation *via* the covalent attachment of ubiquitin. Ubiquitination occurs through three enzymes: a ubiquitin-activating enzyme (E1), a ubiquitin-conjugating enzyme (E2) and s ubiquitin-protein ligase (E3). Upon IFN-γ induction, the key protein degradation machinery is the immunoproteasome, which is a highly complex molecular machine consisting of various complexes, including the 20S core particle. The immunoproteasome 20S core particle has a mass of 700 kDa and comprises 28 protein subunits that are stacked in four homologous rings of seven, forming a hollow cylindrical structure. The two inner rings each formed by seven β subunits (β1i–7i) are enclosed by the two outer rings assembled from seven α subunits (α1–7). The proteolytic chamber is formed by the β rings, which harbor the three catalytically active subunits β1i, β2i and β5i that exhibit caspase-like (CL), trypsin-like (TL), and chymotrypsin-like (ChTL) activities, respectively. Immunoproteasome-degraded products are loaded onto major histocompatibility complex class I (MHC-I) to regulate immune responses by inducing cytotoxic-T-lymphocytes (CTLs). However, when HIV-1 viruses infect the host cells, HIV-1 Nef interacts with LMP7 to attenuate the formation and protein degradation function of immunoproteasome and repress the trafficking and antigen presentation activity of MHC-I.

## Discussion

Immunoproteasome activation plays a crucial role in the immune responses. This study reveals that HIV-1 Nef interacts with LMP7, a key factor for iProteasomes assembly (44), to promote immune evasion through attenuating iProteasome formation. An increasing number of evidences demonstrate important roles of LMP7 in inflammatory diseases. Inflammatory disorders are linked to mutations in LMP7 gene and pharmacological inhibition of LMP7 suggests anti-inflammatory effect in inflammatory models (45, 46). In addition to its function in immune responses, LMP7 plays a role in management of protein homeostasis under oxidative stress and LMP7-deficiency leads to intracellular protein aggregations and autoimmune encephalomyelitis (33, 47).

Formation of iProteasome is a multi-step, delicate, and strictly regulated process, including the assembly of immature 16S precursor, the cleavage of pro-sequences from subunits, and the formation of mature 20S core particle (32, 33). The formation of β ring is on top of assembled α ring, and the adding of β subunits follows a highly strict order (44). Proteasome 20S core particle contains three active subunits with caspase-, trypsin-, and chymotrypsin-like activities that are replaced by related proteases LMP2, MECL1, and LMP7 in iProteasome (48). This exchange of proteolytic activities makes iProteasomes more efficient at generation of peptides for antigen presentation (49). This study demonstrates that Nef interacts with LMP7 on the ER membrane and attenuates incorporation of LMP7 into iProteasome and thereby down-regulates iProteasome formation. Interestingly, both proLMP7-GFP and mLMP7-GFP were able to pull down more Nef than proLMP7 (Fig. 2D), although mLMP7 pulled down less Nef than proLMP7 (Fig. 2B), which may raise a hypothesis that when GFP fused to either full length or mature domain of LMP7, it was able to stabilize their structures and enhance their bindings with Nef. Moreover, Nef represses iProteasome protein degradation function upon IFN stimulation, suggesting that iProteasomes is related to cellular ubiquitination under oxidative stress.

The iProteasome mediates immune responses by efficiently generating peptides for antigen presentation of MHC-I, which is an efficient surveillance system that present antigens when hosts are invaded by pathogens (50). The beginning step in the MHC-I signaling pathway is processing the antigenic proteins into peptides and loading them onto PLCs, in which iProteasomes play a vital role (51). MHC-I molecules present a diverse array of antigenic peptides (immunopeptidome) to circulating TCLs (52). Interestingly, Nef represses the trafficking and antigen presentation activities of MHC-I.

Nef is able to counteract the host immunity (23–27). This study reveals a distinct mechanism by which HIV-1 promotes immune evasion by attenuating the functions of iProteasome and MHC-I through Nef. Nef interacts with LMP7 on the ER membrane and attenuates incorporation of LMP7 into iProteasome to down-regulate the formation and protein degradation function of iProteasome, leading to repress the trafficking and antigen presentation activity of MHC-I.

## Materials and Methods

### Cells and cultures

Human embryotic kidney (HEK) 293T cells, epithelial cervical adenocarcinoma (HeLa) cells and mouse leukaemic monocyte macrophage (RAW264.7) cells were obtained from China Center for Type Culture Collection (CCTCC) (Wuhan, China). Cells were cultured in Dulbecco’s Modification of Eagle’s Medium (DMEM) (Grand Island, NY, USA) containing 5% fetal bovine serum (FBS) (Gibco, Gaithersburg, MD, USA), 1% penicillin, and 1% streptomycin at 37°C with 5% CO_2_. CD4+ Jurkat T cells were obtained from CCTCC and B3Z cell hybridoma was a gift of Dr. Nilabh Shastri of University of California at Berkeley. Cells were cultured in Roswell Park Memorial Institute (RPMI) 1640 medium (Gibco) containing 10% FBS, 1% penicillin, and 1% streptomycin at 37°C with 5% CO_2_. All cell lines were transfected with plasmids using Lipofectamine 2000 (Invitrogen, Carlsbad, CA, USA) following the manufacturer’s instructions.

### Reagents

Recombinant human IFN-γ (rhIFN-γ) was purchased from Peprotech (Rocky Hill, NJ, USA). ER-Tracker Red dyes were purchased from Invitrogen. Egg white ovalbumin (OVA) was purchased from Sangon Biotech (Shanghai, China). NHS-LC-LC-biotin was purchased from Thermo Fisher (Waltham, MA, USA).

### Virus infection

Wild type HIV-1 plasmid (pNL4-3) was obtained from NIH (Rockville, MD, USA) and Nef-deficient HIV-1 plasmid (pNL4-3dNef) was generated from pNL4-3. pNL4-3 and pNL4-3dNef were transfected into HEK293T cells using polyethylenimine (PEI) transfection reagents (Polysciences, PA, USA). At 48 h post-transfection, the cell culture media were collected and the cell debris was removed by centrifugations. Virus titers were measured using p24 enzyme-linked immunosorbent assays (ELISA) (R&D Systems, MN, USA). CD4^+^ Jurkat T cells were incubated with NL4-3 or NL4-3dNef viruses at MOI: 10 ng p24 per 10^6^ cells for 2 h. The cells were washed and replaced with fresh culture media. The cells or supernatants were collected at indicated time for further analyses.

### Stable cell lines

Flag tagged Nef gene was constructed into pLenti vector (Invitrogen). pLenti-Nef plasmids or pLenti empty vectors were co-transfected with pLP1, pLP2, and pLP/VSVG plasmids (Invitrogen) into HEK293T cells by using PEI transfection reagents. The cell supernatants containing Lentiviruses were collected after 2 days of transfections. CD4^+^ Jurkat T cells were infected with Lentiviruses containing Nef or empty vector respectively. Stable Nef expressing cells were selected with 2.5 µg/ml puromycin.

### Protein polyubiquitylation

HeLa cells were incubated with 100 U/ml rhIFN-γ after 1 day of transfection with Nef plasmids or empty vectors. Alternatively, CD4^+^ Jurkat cells were incubated with 100 U/ml rhIFN-γ after 2 h of infections with wild type NL4-3 or Nef deficient NL4-3dNef viruses. The cells were harvested and lysed for western blots at indicated time.

### Plasmids construction

The full length and truncations of Nef were constructed into pCAGGS vectors (BCCM/LMBP Plasmid Collection, Gent, Belgium) with an HA tag linked to the C terminal insert. Nef mutants were also constructed into pCAGGS vectors using overlapping polymerase chain reaction (PCR), and the putative amino acid residues were mutated to Alanine. Alternatively, the full length of Nef was constructed into the pLenti vector (Invitrogen) with a green fluorescent protein (GFP) gene fused to the N terminus of the Nef gene. The full length and truncations of LMP7 were constructed into pcDNA3.1 vectors (Invitrogen) with a 3 x Flag tag linked to the N terminal insert. Alternatively, the full length LMP7 was constructed into the pCMV vector (Clontech, Fremont, CA, USA) with a Myc tag linked to the N terminal insert. Nef was constructed into the pGBKT7 vector and LMP7 was constructed into the pGADT7 vector (Clontech) in the yeast two-hybrid assays. The XhoI restriction enzyme site in pNL4-3 was digested and inserted a stop codon to generate pNL4-3dNef.

### Confocal microscopy

HeLa cells were seeded in 15 mm glass bottom dishes (Nest Scientific, NJ, USA) 1 day before transfections. 1 µg plasmids were transfected for each dish. Cells were washed 3 times with phosphate buffer saline (PBS) and fixed with 1% formaldehyde in PBS for 30 min at 24 h post transfections. After 3 times washes with PBS, cells were permeabilized in buffer (PBS containing 1% bovine serum albumin and 0.1% saponin) for 30 min, and then incubated with anti-HA or anti-Flag (Sigma-Aldrich, MO, USA) for 1 h. Cells were washed 3 times with permeabilization buffer, and then incubated with secondary antibodies for 1 h. After 3 times washes with PBS, cells were stained with endoplasmic reticulum (ER)-Tracker Red dyes (Invitrogen) for 30 min. Then cells were washes 3 times in PBS and viewed by using FluoView FV1000 (Olympus, Tokyo, Japan) confocal microscope.

### Recombinant protein expression and purification

Nef and LMP7 genes were cloned into pGEX-6P-1 (GE, Boston, MA, USA) and pET-28a plasmids respectively (Novagen, Madison, WI, USA). pGEX-6P-1 empty plasmid and pEGX-6P-1-Nef plasmid were respectively transformed into BL21 (DE3) competent cells (TransGen, Beijing, China). Alternatively, pET-28a-LMP7 plasmids were transformed into BL21 (DE3) competent cells (TransGen). The cells were grown in LB buffer with 0.1mM IPTG at 30°C for 16 h, then harvested by centrifugation at 3,000 g for 10 min. The cell pellets were re-suspended in PBS, then frozen and thawed for 3 times, and sonicated for 10 min (pulse on for 5 s and pulse off for 5 s) on ice. The supernatants were collected by centrifugation at 12,000 rpm for 10 min, then the proteins were purified by using NGC Scout10 (Bio-Rad, Hercules, CA, USA) according to manufacturer’s protocol.

### In vitro GST pull down assay

LMP7 proteins expressed in bacterial cells, and were mixed with purified glutathione-S-transferase (GST) or GST-Nef proteins in RIPA buffer. The samples were mixed with recombinant protein G-Sepharose 4B beads (Thermo Fisher) and 1 µl anti-Flag antibodies (Sigma-Aldrich) at 4°C for overnight. The beads were washed with RIPA buffer for 4 times, and mixed with loading buffer, then run on sodium dodecyl sulfate-polyacrylamide gel electrophoresis (SDS-PAGE) gels and subjected to western blots.

### Biolayer interferometry (BLI) assay

Purified LMP7 proteins from bacteria were linked with NHS-LC-LC-biotin (Thermo Fisher) as a 1:3 molar ratio at room temperature for 1 h. Proteins were clarified through Zeba spin desalting columns (Thermo Fisher) to remove un-reacted biotin. The Streptavidin biosensors (ForteBio, Menlo Park, CA, USA) were sequentially dipped into wells of black 96-well plates (Greiner, Kremsmünster, Austria) containing PBS, LMP7, PBS, GST or Nef-GST, and PBS. All the association and dissociation curves were fitted 1:1 binding model.

### Proteasome purification

HeLa cells were seeded in 10-cm dishes 1 day before transfections. 10 µg Nef-HA or empty vector plasmids were transfected into the cells at 50–80% confluence for each dish by mixing with 20 µl PEI transfection reagents. After 24 h post transfection, the cells were treated with 100 U/ml IFN-γ (Peprotech, Rocky Hill, NJ, USA) for 24 h. Cells were trypsinized after washed with pre-chilled PBS, then lysed in 1 ml buffer A (50 mM Tris-HCl, pH7.5, 250 mM sucrose, 150 mM NaCl) by 3 times of freeze and thaw for each dish. Cell lysates were pre-clarified by centrifugation at 1,000 g for 5 min at 4°C. The supernatant was collected and clarified again at 10,000 g for 20 min at 4°C. A 20 µl supernatant aliquot of each sample was saved and stored at 4°C until further analysis. The rest of the samples were spun in buffer A at 100,000 g for 1 h at 16°C. After discarding the pellets, the supernatant was spun at 100,000 g for 5 h, at 16°C. The pellets were dissolved in 20 µl buffer B (50 mM Tris-HCl, pH7.5, 150 mM NaCl, 15% glycerol) for each sample then subjected to western blots for further analysis.

### Western blot and immunoprecipitation

HEK293T cells were plated in 6 cm dishes 1 day before transfections. A total amount of 4 µg plasmids was transfected for each plate by mixing with 8 µl PEI transfection reagents. At 24 h post-transfections, cells were lysed in radio immunoprecipitation assay (RIPA) buffer (50 mM Tris-HCl, pH7.4, 150 mM NaCl, 0.25% deoxycholate, 1% NP-40) with 1 mM phenylmethanesulfonyl fluoride (PMSF) (Sigma-Aldrich). Cell lysate supernatants were clarified by centrifugation at 12,000 rpm for 10 min, then mixed with SDS-PAGE loading buffer (50 mM Tris-HCl, pH6.8, 2% SDS, 10% glycerol, 0.1% bromophenol blue, 1% 2-mercaptoethanol), and heated at 95°C for 10 min. Supernatants were loaded in 12% SDS-PAGE gels and run at 120 V for 2 h. The proteins were transferred to polyvinylidene difluoride (PVDF) membranes (GE) at 70 V for 2 h. Membranes were blocked with 2% nonfat milk in phosphate buffer solution (PBST) buffer (0.1% tween-20 in PBS) for 30 min at room temperature. After washed 3 times with PBST, membranes were incubated with primary antibodies overnight. The membranes were then incubated with secondary antibodies for 1 h after washed 3 times with PBST. Following 6 times washes of PBST, the membranes were incubated with clarity western ECL substrate (Bio-Rad), then scanned by LAS-4000 (FujiFilm, Tokyo, Japan). Alternatively, cell lysate supernatants were mixed with antibodies and recombinant protein G-Sepharose 4B beads (Thermo Fisher) at 4°C for overnight. The beads were washed 4 times with RIPA buffer, then mixed with SDS-PAGE loading buffer and run on the SDS-PAGE gels.

### Flow cytometry

CD4^+^ Jurkat T cells stably expressing Nef-Flag or empty vector were seeded in 6-well plates 1 day before transfections. 50 nM LMP7 or control siRNA was transfected into the cells by mixing with 7 µl INTERFERin transfection reagents (Polyplus-transfection, Illkirch, France) for each well. After 24 h transfections of siRNA, cells were treated with 100 U/ml rhIFN-γ for 24 h. The cells were collected and incubated with block reagents (1% bovine serum albumin, BSA in PBS) (Sigma-Aldrich) for 30 min after fixing with 1% formaldehyde in PBS for 10 min. The cells were incubated with W6/32 APC or isotype control antibodies for 1 h. After filtering with membranes, the cells were analyzed by using Cytoflex (Beckman, Brea, CA, USA), and 50,000 events were collected for each sample.

### Antigen presentation assay

RAW264.7 cells were seeded in 24-well plates. After cells attached to the plates, 20 ng purified GST or GST-Nef proteins purified from bacteria were added into the cell media for each well at 50% cell confluence for 15 h. After 3 times washes with PBS, cell media was replaced by Opti-MEM I reduced serum medium (Thermo Fisher) with 10 mg/ml egg white ovalbumin (OVA) (Sangon Biotech, Shanghai, China) for 6 h. After 3 times washes with PBS, the cells were co-cultured with B3Z cells for 24 h. The cell culture media was collected and spun at 12,000 rpm for 10 min to discard the cell debris then frozen at −80°C until further measurements. ELISA was performed to measure IL-2 concentration of the thawed samples by using mouse IL-2 ELISA set (BD Biosciences, Franklin Lakes, NJ, USA) according to manufacturer’s instructions.

### Yeast two-hybrid assay

Nef and LMP7 genes were constructed into pGBKT7 and pGADT7 vectors (Clontech), respectively. pGBKT7-Nef and pGADT7-LMP7 plasmids were transformed into Y2HGold and Y187 yeast strains to grow at 30°C for 3 days. Y2HGold and Y187 yeasts (Clontech) were picked and mix together to grow in YPDA broth at 200 rpm, 30°C for overnight. The mated culture was plated on DDO (-Leu/-Trp) agar plates to grow at 30°C for 3 days. Colonies were picked and streaked on QDO (-Ade/-His/-Leu/-Trp), and X-α-Gal agar plates to grow at 30°C for 3 days.

### Statistics

All experiments were repeated at least three times with similar results. All results were expressed as the mean ± the standard deviation (SD). Statistical analysis was carried out using the two-tailed t-test (GraphPad Prism5). The date difference was considered statistically significant when *P* ≤0.05 (*), *P* ≤0.01 (**), *P* ≤0.001 (***); NS, not significant.

## Acknowledgments

We thank Dr. Nilabh Shastri of University of California at Berkeley for kindly providing the T-cell hybridoma cell line B3Z.

This work was supported by National Natural Science Foundation of China (81730061 and 31800147), National Health and Family Planning Commission of the People’s Republic of China, National Mega Project on Major Infectious Disease Prevention (2017ZX10103005 and 2017ZX10202201), Guangdong Province “Pearl River Talent Plan” Innovation and Entrepreneurship Team Project (2017ZT07Y580).

The authors declare that they have no competing interests.

